# Gene Regulatory Network Inference as Relaxed Graph Matching

**DOI:** 10.1101/2020.06.23.167999

**Authors:** Deborah Weighill, Marouen Ben Guebila, Camila Lopes-Ramos, Kimberly Glass, John Quackenbush, John Platig, Rebekka Burkholz

## Abstract

Gene regulatory network inference is instrumental to the discovery of genetic mechanisms driving diverse diseases, including cancer. Here, we present a theoretical framework for PANDA, an established method for gene regulatory network inference. PANDA is based on iterative message passing updates that resemble the gradient descent of an optimization problem, OTTER, which can be interpreted as relaxed inexact graph matching between a gene-gene co-expression and a protein-protein interaction matrix. The solutions of OTTER can be derived explicitly and inspire an alternative spectral algorithm, for which we can provide network recovery guarantees. We compare different solution approaches of OTTER to other inference methods using three biological data sets, which we make publicly available to offer a new application venue for relaxed graph matching in gene regulatory network inference. We find that using modern gradient descent methods with superior convergence properties solving OTTER outperforms state-of-the-art gene regulatory network inference methods in predicting binding of transcription factors to regulatory regions.

## 1 Introduction

Next generation genome sequencing technology has revolutionized genetic research and provides data at an unprecedented scale. This progress facilitates large, genome-scale studies which provide new insights into gene regulation, including the control of protein production through the expression of genes. Proteins influence higher level cellular functions, which are often altered during the development and progression of different diseases, including cancer. To gain an understanding of the gene regulatory mechanisms perturbed by a disease, it is common practice to infer and compare associated gene regulatory networks (GRNs) [20, 22, 26, 34]. In many cases, these networks are weighted, bipartite, and have a representation as matrix *W*. *W* consist of two types of nodes – transcription factors (TFs) and genes. A TF is a protein that can bind to the DNA in the vicinity of a gene and regulate its expression, which constitutes a link in the gene regulatory network, see Fig. 1. Gene expression often leads to the production of associated proteins (including TFs) that can interact, form higher-order protein complexes, regulate genes, etc. Possible interactions of such proteins are studied in detail and are commonly summarized in a known matrix *P*. Also data on gene expression is widely accessible but is context specific, i.e. it depends on the tissue type, disease, etc. Based on this data, we can define a gene-gene co-expression matrix *C*. A more detailed explanation of gene regulation is given in the supplement. We also recommend the review article by Todeschini et al. [30]).

**Figure 1:**
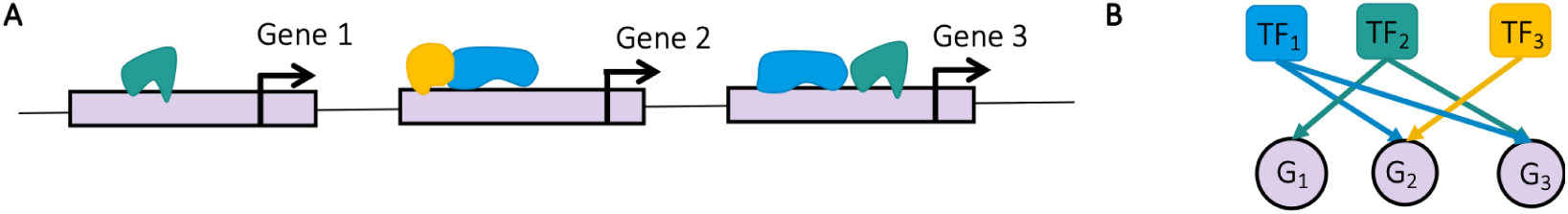
Gene regulation. A. Transcription factors (TFs) are represented by green, blue, and yellow objects that bind to the genome (gray band) in vicinity of the start site of a gene (black arrow) to regulate its expression. B. Representation of A as bipartite gene regulatory network.

Our main goal is to infer a gene regulatory matrix *W* given the two matrices *P* and *C*. We pose this as a non-convex optimization problem, OTTER (**O**p**t**imize **t**o **E**stimate **R**egulation). It is related to relaxed inexact graph matching, which seeks agreement between two graphs (that are represented by matrices like *P* and *C*). The gene regulatory matrix *W* could be seen as relaxed permutation matrix that matches vertices in *P* and *C*. Similarly to graph matching, OTTER is theoretically tractable but the solutions are nonunique, which explains in part why GRN inference is challenging. We characterize the solution space and propose two distinct approaches for model selection, a spectral approach and gradient descent. For both, we provide theoretical network recovery guarantees. While the spectral method is robust to small noise, gradient descent is more reliable in higher noise settings and outperforms state-of-the-art gene regulatory network inference techniques.

### Contributions

1) We pose a novel optimization problem for gene regulatory network inference, OTTER, which is analytically tractable. 2) We gain insights into a state-of-the-art GRN inference method, PANDA [12], as OTTER gradient descent resembles the related message passing equations. 3) We characterize OTTER’s solution space and derive a spectral algorithm on this basis, for which we give network recovery guarantees. 4) We solve the gradient flow dynamics associated with gradient descent for OTTER. 5) We draw a connection from OTTER to relaxed graph matching and open a new application area for related algorithms. 6) OTTER gradient descent outperforms the current state of the art in GRN inference on three challenging biological data sets related to cancer. 7) We make the processed data publicly available to ease the use for researchers without a genetics background and to foster furthter innovation in relaxed graph matching and GRN inference.

### Related work

The OTTER objective is inspired by a state-of-the-art GRN inference method, PANDA (Passing Attributes between Networks for Data Assimilation) [12]. PANDA integrates multiple data sources through a message passing approach, which we find resembles the gradient descent of OTTER. A derivation is given in the supplement. PANDA has been used to investigate gene regulatory relationships in both tissue specific [28] as well as several disease contexts, including chronic obstructive pulmonary disease [20], asthma [26], beta cell differentiation [34], and colon cancer [22]. OTTER can be seen as a theoretically tractable simplification of PANDA, which is amenable to modern optimization techniques and draws connections to graph matching.

Graph matching [32] has strong theoretical foundations [16, 3] that can benefit a deeper understanding of OTTER and vice versa. In particular, the quadratic assignment problem (QAP) [1] and its variants [24] have a direct link to OTTER and can support similar biological theory. Graph matching has broad applications in computer science ranging from machine learning [7], pattern matching [36], vision [4, 35], and protein network alignment [27] to social network analysis [8], but has not been applied to gene regulatory network inference to the best of our knowledge. As we show, simple relaxed graph matching techniques outperform established GRN inference methods, which are detailed in the experiment section (Sec. 4).

## 2 Theoretical framework

Our goal is to learn a bipartite and weighted gene regulatory network which we represent as a matrix 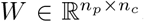. An entry *w*_*ij*_ indicates whether TF protein *i* binds to a region of the DNA that is associated with gene *j* and has evidence that it regulates that gene. A larger weight *w*_*ij*_ is assumed to be associated with a higher probability of binding. We have *n*_*p*_ TFs and *n*_*c*_ genes, where the number of genes *n*_*c*_ is much larger (*n*_*p*_ ≪ *n*_*c*_). We assume that we observe only squares of *W*, which are given as symmetric matrices 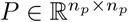 (the protein-protein interaction matrix) and 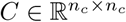 (the gene-gene co-expression matrix). Thus, *W* explains protein interactions as *WW*^*T*^ ≈ *P* and gene co-expression as *W*^*T*^*W* ≈ *C*. We formulate this reasoning as the following optimization problem:

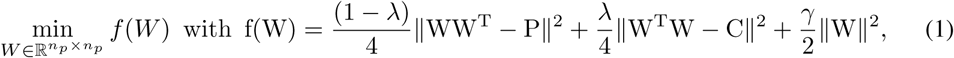

which we call OTTER. *λ* ∈ [0, 1] denotes a tuning parameter that moderates the influence of *P* versus *C*, and *γ* corresponds to a potential regularization. In principle, we could choose any matrix norm but limit our following discussion to the Frobenius norm 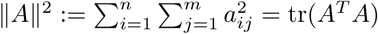 for a matrix *A* = (*a*_*ij*_). For this choice, gradient descent resembles most closely the related message passing equations of PANDA and we can derive the corresponding solutions.

These solutions depend on the spectral decomposition of 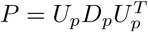 and 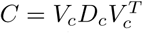, which exist with respect to orthogonal *U*_*p*_ and *V*_*c*_, as *P* and *C* are symmetric. Otherwise, the same results hold for the spectral decomposition of (*P* + *P*^*T*^)*/*2 and (*C* + *C*^*T*^)*/*2. *D*_*p*_ and *D*_*c*_ are diagonal matrices containing the eigenvalues of the respective matrix. In a slight abuse of notation, we denote with *D*_*p*_ a matrix 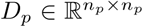 and, if convenient, a matrix 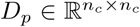, which is padded with zeros accordingly. Furthermore, let 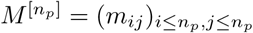 denote a submatrix of a larger matrix *M* with dimension *n*_*p*_ × *n*_*p*_. Without loss of generality, we assume that the eigenvalues *d*_*p,ii*_ of *P* are indexed in descending order; *d*_*p,ii*_ ≥ *d*_*p,jj*_ for *i < j*. For *C* however, we require a good matching with *P*. We therefore assume implicitly that the distance of *D*_*c*_ to *D*_*p*_ is minimized with respect to permutations of the eigenvalues of *C*, that is ‖*D*_*c*_ − *D*_*p*_‖^2^ = min_*π*∈𝒫_‖*D*_*c,π*_ − *D*_*p*_‖^2^, where 𝒫 denotes the set of permutations of {1, …, *n*_*c*_} and *D*_*c,π*_ the corresponding ordering of eigenvalues on the diagonal. If *D*_*p*_ and *D*_*c*_ show little discrepancy, this will result in the eigenvalues of *C* being in descending order as well. Note that 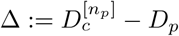 measures the discrepancy in our biological hypothesis from the start. Usually, we can further assume that *P* and *C* are positive semi-definite so that *d*_*ii*_ ≥ 0, meaning that our model is specified well enough such that *C* and *P* have roots. Based on this, we can characterize the solution space 𝒮 in the general case.

### Theorem 1.

*For given* 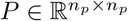 *with P* = *P*^*T*^ *and* 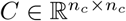 *with C* = *C*^*T*^, *for any spectral decomposition* 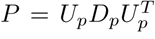 *and* 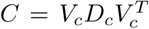, *λ* ∈ [0, 1], *the minimization problem (1) has solutions W* ^∗^ ∈ 𝒮 *with singular value decomposition* 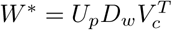, *where*

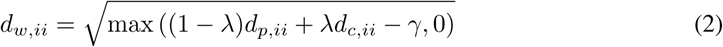

*for i* ≤ *n*_*p*_. *For d*_*w,ii*_ = 0, *the corresponding columns of U*_*w*_ *and V*_*w*_ *are not restricted to the eigenvectors of P and C. The eigenvalues of C are ordered such that D*_*c*_ = *D*_*c,π*_, *where the permutation solves the minimization problem*

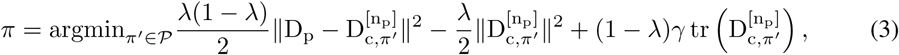

*For* 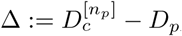, *we further assume that*

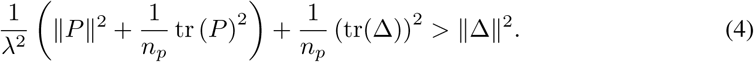

Condition (4) is usually met and it is a minor technicality to exclude alternative global minima of Objective (1) that defy our intuition. The nature of these alternatives is discussed in detail in the proof of the Theorem in the supplement.

According to Thm. 1, OTTER (1) has at least 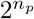 different solutions. Each column *u*_:*i*_ of *U*_*p*_ has two optional signs that do not alter the spectral decomposition of *P* but can lead to a different *W* ^∗^. The same applies to columns *v*_:*i*_ of *V*_*c*_. Only the product of corresponding columns (*u*_:*i*_ and *v*_:,*i*_) determines the respective solution *W* ^∗^, as we have 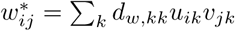. This leaves us with 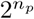 alternatives. If the spectra are not simple, such that some eigenspaces have multiple choices of basis functions, we have additional degrees of freedom in constructing the solutions.

As a consequence, we face a model selection problem and require additional information to make an informed decision. Indeed, PANDA takes as an additional input an initial guess of a gene regulatory matrix 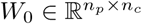, which is typically based on data from a scan of the genome sequence using known transcription factor binding motifs. Assuming that *W*_0_ provides good evidence, our first proposal selects the closest solution to *W*_0_ using a spectral approach.

### 2.1 A spectral method for solving OTTER

For additional evidence *W*_0_, we turn problem (1) into

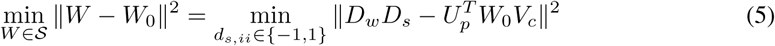

where we fix *U*_*p*_ and *V*_*c*_ and assume that *W* has simple singular values, meaning the singular values are different from each other. For simplicity, we write 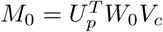. For simple spectra of *P* and *C* and matched eigenvalues, *W* ^∗^ is unique and given by 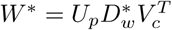 with 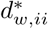 = *d*_*w,ii*_sign(m_0,ii_), where *d*_*w,ii*_ is defined as in Thm. 1 and sign(x) = 1 for *x* ≥ 0 and sign(x) = −1 for *x <* 0.

The question is how well this approach performs in the presence of noise. As the next proposition shows, even if only *W*_0_ is noise corrupted so that we know *D*_*W*_, perfect recovery of *W* is unlikely for large-scale problems. Let Φ denote the cumulative distribution function (cdf) of a standard normal and *X* ∼ Ber(p) a Bernoulli random variable with success probability *p*.

#### Proposition 2.

*Assume that we observe P* = *W* ^∗*T*^*W* ^∗^, *C* = *W* ^∗^*W* ^∗*T*^, *and W*_0_ = *W* ^∗^ + *ϵ for a true underlying* 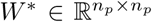 *and noise* 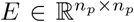 *with independent identically normally distributed components e*_*ij*_ ∼ 𝒩 (0, *σ*^2^). *Further assume that P and C have a simple spectrum* 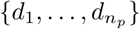. *Then, for the spectral approach* 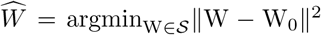 *with γ* = 0, *the recovery loss is distributed as* 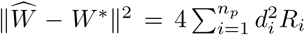, *where R*_*i*_ ∼ Ber (Φ (−d_i_*/σ*)) *for d*_*i*_ > 0 *and R*_*i*_ = 0 *for d*_*i*_ = 0 *are independent. For any ϵ* > 0, *the following holds with the usual Chernoff bound:*

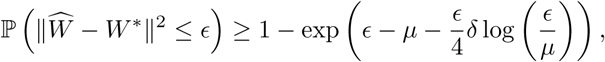

*where µ* = ∑_*i*_ *p*_*i*_ *and* 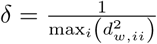 *for ϵ* ≤ *µ and* 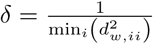 *otherwise.*

In particular, for the probability of perfect recovery (*ϵ* = 0) we have 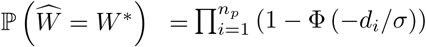. In our examples, *n*_*p*_ = 1636 and *d*_min_ ≈ 0.0001. To achieve a probability of at least 0.5, we could allow for a noise variance of *σ* ≈ 3 · 10^−5^. In many applications, this would be a reasonable range, considering that we have *n*_*p*_*n*_*c*_ ≈ 4.4 · 10^7^ matrix entries. Biological data is known to be noisy, so we expect levels of noise in *P* and *C* that will require regularization.

#### Regularization

Accounting for noise in *P* and *C* complicates the analysis considerably, as the spectral decomposition of *P* and *C* (including their eigenvectors) are distorted. We focus on the population version (the average matrices) to motivate the need for additional regularization (*γ* > 0). Depending on the source of the noise, the spectrum can become biased. To see this, assume noise of the form *P* = (*W* ^∗^ + *E*_*p*_)(*W* ^∗^ + *E*_*p*_)^*T*^ and *C* = (*W* ^∗^ + *E*_*c*_)^*T*^ (*W* ^∗^ + *E*_*c*_) or 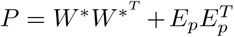 and 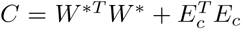, respectively. If *E*_*p*_ and *E*_*c*_ have iid entries with zero mean, variance 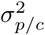, and a symmetric distribution, we get 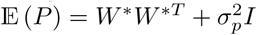 and 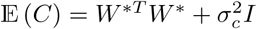, where *I* denotes the respective identity matrix. The choice 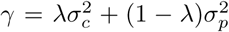 in Eq. (1) can compensate for this spectral shift. It should be noted that, Thm. 1 states that such a l2 regularization alters the solutions to Problem (1) in two ways. Not only are the singular values of *W* ^∗^ shifted by −*γ* to compensate for the biases introduced by the noise, also the matching of the eigenvalues of *P* and *C* is influenced by the additional penalty 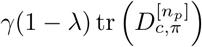 in Eq. (3). Consequently, it may be optimal to pair the eigenvalues of *P* with smaller eigenvalues of *C* rather than larger ones if *γ* is large.

While the spectral method can be powerful in a setting in which noise is controlled such that our assumptions are met approximately, the gradient descent alternative gives us more tuning options, including the step size and early stopping, that will allow us to stay closer to the initial guess *W*_0_.

### 2.2 Gradient descent for solving OTTER

Indeed, the message passing equations of PANDA resemble a gradient descent procedure, where the gradient of Objective (1) is given as ∇ *f* (*W*) = *WW*^*T*^*W* − (1 − *λ*)*PW* − *λWC* + *γW*. We explain this relationship in detail in the supplement. In our experiments, we used the ADAM method [18] for gradient descent but alternatives are equally applicable. From a theoretical perspective, we can understand the gradient dynamics for specific choices of *W*_0_. For this purpose, we take the continuous time approximation (corresponding to infinitesimally small step size) and study the corresponding gradient flow:

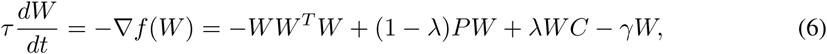

where we set the time unit *τ* = 1 in the following for simplicity. If the initial *W*_0_ has a similar singular value decomposition as a solution, the differential equation decouples and we can solve the resulting one-dimensional ordinary differential equations for the diagonal elements explicitly.

#### Proposition 3.

*For initial* 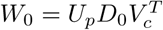 *with* 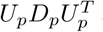 *and* 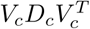, *the solution of the gradient flow (6) is given by* 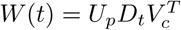 *with*

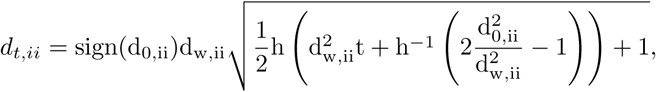

*where h*(*x*) = tanh(x) *if* 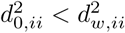 *and h*(*x*) = coth(x) *otherwise.*

Note that the square root factor converges to 1 for *t* → ∞ in both cases. Hence, the final solution inherits the signs sign(d_0,kk_) of the initialization similar to our spectral approach. Thus, if we start from a reasonable guess *W*_0_ that diagonalizes with respect to the same *U* and *V* as the global minima, gradient descent will converge to the closest global minimum for small enough learning rate. For general *W*_0_, however, it is important to keep in mind that gradient descent can converge to different solutions. It does not necessarily stay close to our initialization and can even get stuck in a local minimum.

## 3 Relation to inexact graph matching

The biological justification for the OTTER objective function is also consistent with quadratic assignments [1], a more common choice in graph matching. From *P* ≈ *WW*^*T*^ and *C* ≈ *W*^*T*^*W* we could deduce *PW* ≈ *WW*^*T*^*W* ≈ *WC* and thus minimize the Quadratic Assignment Problem (QAP) objective

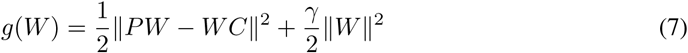

(with additional l2-regularization). In graph matching however, *P* and *C* are usually assumed to have the same dimension. They differ for inexact graph matching, but the smaller network is then supposed to be similar to a subgraph of the bigger one. Thus, the minimization is performed under the constraint that *W* is a permutation matrix. In contrast, we are interested not in a permutation matrix, but in a weighted network 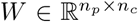 that solves the relaxed QAP. The corresponding gradient is ∇*g*(*W*) = *P* ^2^*W* + *WC*^2^ − 2*PWC* + *γW*. While the related ADAM gradient descent approach is computationally more costly than the one solving OTTER, both approaches perform similarly well as we show in experiments (see Sec. 4).

This indicates that there is a great potential to perform gene regulatory network inference using alternative graph matching methods in future investigations. In particular, GRAMPA [9, 10] is a variant of QAP with strong recovery guarantees, also for the Wigner model where *P* − *C* has iid noise entries. GRAMPA adds the term −*δ***1**^*T*^*W* **1** to the QAP objective (7), where **1** denotes a vector with all entries equal to one. As a consequence, the solution to the minimization problem becomes unique and explicitly solvable using a spectral approach. As the spectral version of OTTER, it does not perform well in estimating GRNs. Solving the GRAMPA objective function with gradient descent works better, though it is not competitive.

Graph matching can also be studied within the optimal transport framework [25, 29]. We could formulate the OTTER objective with respect to a nonstandard metric and regularization term. Since we are not searching for stochastic matrices *W*, this does not serve our purpose and we leave the transfer of related methods to gene regulatory network inference to future explorations.

## 4 Experiments

### 4.1 Experiments on synthetic data

To showcase the performance of OTTER for cases in which our assumptions are met, we create artificial data based on a ground truth *W* ^∗^ that we try to recover from noise corrupted inputs *W*_0_ = *W* ^∗^ + *N*_0_, *P* = (*W* ^∗^ + *N*_*p*_)(*W* ^∗^ + *N*_*p*_)^*T*^, and *C* = (*W* ^∗^ + *N*_*c*_)^*T*^ (*W* ^∗^ + *N*_*c*_). All noise entries are Gaussian and independently distributed with 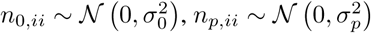, and 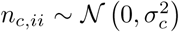. To obtain a realistic ground truth for which we can repeat each experiment 10 times conveniently, we sub-sample (in each repetition) the ChIP-Seq network for the liver tissue to *n*_*p*_ = 100 and *n*_*c*_ = 200. (See the next section for more details.) As this is unweighted, we draw the weights iid from 𝒩(10^−5^, 1). For each network, we use the spectral and the gradient descent version of OTTER and report the obtained recovery error ‖*W* − *W* ^∗^‖^2^.

For simplicity, we do not align the eigenvalues of *P* and *C* optimally but arrange each of them in descending order in the spectral approach. We run ADAM gradient descent for the OTTER objective for 10^4^ steps with the default ADAM parameters, as detailed in the supplement. For both the gradient decent and the spectral approach, we use parameters 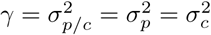 and *λ* = 0.5.

The results are shown in Fig. 2. For small levels of noise in *P* and *C*, the spectral approach performs reliably and better than gradient descent. However, for high 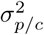 gradient descent outperforms the spectral method. Since we biological data are inherently noisy, gradient descent seems to be the method of choice. Furthermore, it provides us with additional tuning options that we can leverage to outperform state of the art methods.

**Figure 2:**
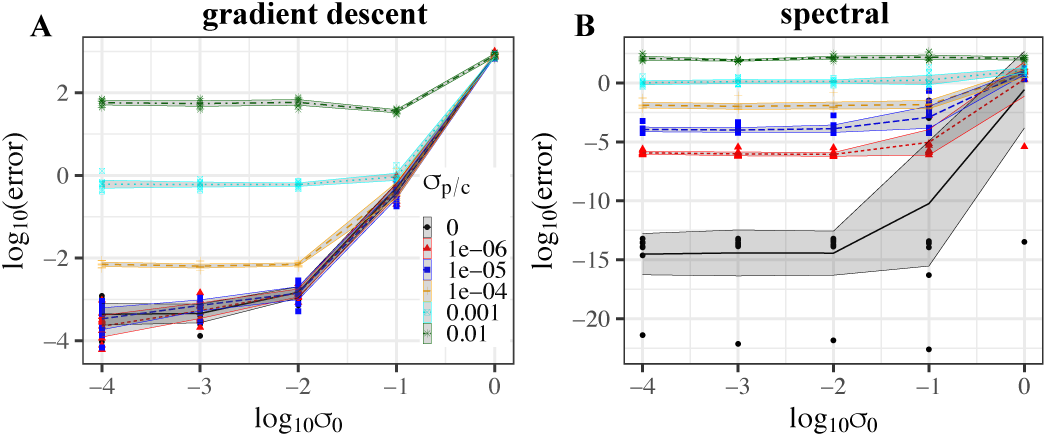
OTTER recovery error using (A) gradient descent and (B) spectral decomposition for artificial networks of size *n*_*p*_ = 100, *n*_*c*_ = 200 and Gaussian noise with variance 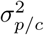 for *P* and *C* and 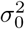 for *W*_0_. Shaded regions correspond to the 0.95 confidence interval and lines to the average over 10 repetitions. The legend applies to both figures.

### 4.2 Experiments on cancer data

The most abundant data source for studying gene regulation is gene expression data. These data are often measured using bulk RNA-sequencing (RNA-seq) with samples corresponding to different individuals.

#### Datasets and experimental set-up

We obtained bulk RNA-seq data from the Cancer Genome Atlas (TCGA) [31]. The data is downloaded from recount2 [6] for liver, cervical, and breast cancer tumors and normalized and filtered as described in the supplement. The corresponding Pearson correlation matrix defines the gene-gene co-expression matrix *C* consisting of *n*_*c*_ = 31, 247 genes for breast cancer, *n*_*c*_ = 30, 181 for cervix cancer and 27, 081 for liver cancer. The protein-protein interaction matrix *P* is derived using laboratory experiments and represents possible interactions; we use a padded version of that provided in [28]. It consists of *n*_*p*_ = 1, 636 potential TFs. Our initial guess of a gene regulatory network, *W*_0_, is similar across tissues but varies slightly depending on the number of genes (*n*_*c*_) included after filtering and normalization. *W*_0_ is a binary matrix where there is a “1” if there is a TF sequence motif in the promoter of the target gene, and a “0” otherwise. Sequence motif mapping was performed using the FIMO software [13] from the MEME suite [2] and the GenomicRanges R package [21]. In addition to the standard PANDA input *P* and *W*_0_, we also consider transformations thereof, i.e., 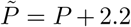 with element-wise addition and 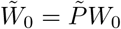, that have been inspired by aspects of GRAMPA graph matching and increase the performance of most methods considerably. Note that neither *W*_0_, 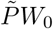 or *P* carry sign information about edge weights so that we cannot infer whether TFs inhibit or activate the expression of a gene. We therefore focus on the prediction of link existence with the understanding that the type of interaction can be estimated *post hoc*.

Validation of gene regulatory networks is a major challenge. Data from Chromatin immunoprecipitation (ChIP)-sequencing (ChIP-seq) experiments, which measures the binding of TFs to DNA in the genome, provides a validation standard against which to benchmark our results. It provides only partial validation as only a small fraction or the collections of TFs are typically profiled for any tissue.

However, because of the destructive nature of collecting molecular data from biological samples, ChIP-seq and RNA-seq data cannot be easily collected from the same biological samples, and so ChIP-seq data is generally collected from cell lines in the laboratory rather than from patient tumor samples. We used ChIP-seq data from the HeLa cell line (cervical cancer, 48 TFs), HepG2 cell line (liver cancer, 77 TFs) and MCF7 cell line (breast cancer, 62 TFs) available in the ReMap2018 database [5]. Because of the limitations in the available data we use AUC-ROC (area under the receiver operating characteristic curve) and AUPR (or AUC-PR) (area under the precision recall curve) to measure the performance of link classification on the subnetwork in each tissue that is constrained to the measured TFs.

Hyperparameter tuning of OTTER was assisted by MATLAB’s bayesopt function utilizing a Gaussian process prior to maximize the joint AUC-PR for breast and cervix cancer, max AUPR_breast_ · AUPR_cervix_. Breast and cervix data serve therefore as training data while the liver cancer data is an independent test set. The parameters of all compared methods are reported in the supplementary information.

#### Related literature and methods

Many methods try to infer regulatory relationships solely based on gene expression with two possible (non-exclusive) objectives: structure learning and gene expression prediction. Note that the latter usually includes the former. TFs are proteins that are created from the mRNA expressed by their corresponding genes. Hence, predicting target gene expression from the expression of the genes coding for the TFs assumes a biologically reasonable structure.

The most common and basic approach is to analyse the Pearson correlation (COR) matrix or, if feasible, partial correlations (PARTIAL COR). Alternatives are based on mutual information, where ARACNe [19] is one of the most commonly used representatives. Among graphical models, mainly Gaussian graphical models are used because the learning algorithms have to scale to approximately *n*_*c*_ = 30000 genes (in the case of human tissue). The GLASSO [11] method is among the best performing candidates and uses LASSO regularization to enforce sparsity. However, it still does not scale to our setting, and thus we have omitted it from our analysis. Linear models [14] and random forests [15] have been used for a similar purpose, where TIGRESS [14] and GENIE3 [15] were top scorers at the DREAM5 challenge [23] (although the challenge was somewhat different from the GRN modeling we describe here). Both methods have high computational requirements and are less suitable for the human genome that consists of more than 25,000 genes. An alternative approach is to treat ChIP-seq binding predictions as a supervised learning problem [17, 33]. While such models can be quite accurate, they are limited to the small number of TFs for which ChIP-seq experiments are available and thus limited in their discovery of new gene regulatory relationships. Note that, in contrast, *P* and *W*_0_ for PANDA-related objectives are always available and include a much larger set of known TFs.

#### Results

Table 1 compares the feasible GRN inference and relaxed graph matching methods based on comparison to experimental ChiP-seq binding data. Overall, the OTTER^∗^ (or OTTER) gradient descent approach achieves the best performance on all tissues, in particular, on the liver test set (see also Fig. 3). An enrichment analysis of Gene Ontology terms between networks for healthy and cancerous liver tissue in the supplement provides additional evidence that OTTER^∗^ (GRAD) is biologically meaningful. The ADAM gradient descent solving the relaxed quadratic assignment problem (QAP^∗^ grad) is almost on par with OTTER^∗^ (GRAD).

**Table 1:**
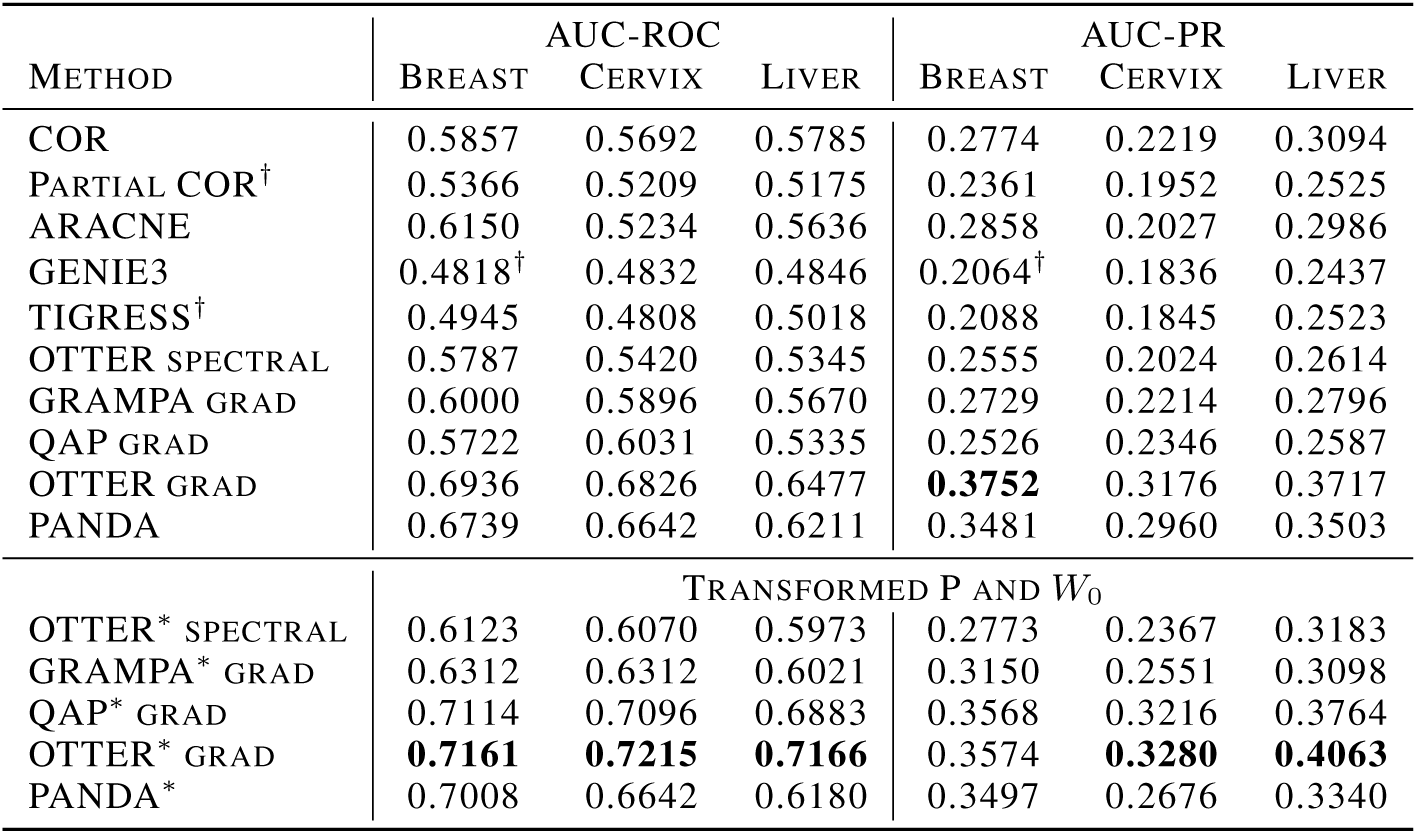
TF binding prediction for different cancer tissues. The symbol † indicates that binding predictions were made only for TFs with ChIP-seq data due to high computational demands. The highest AUC-ROC/AUC-PR for each data set is shown in bold.

**Figure 3:**
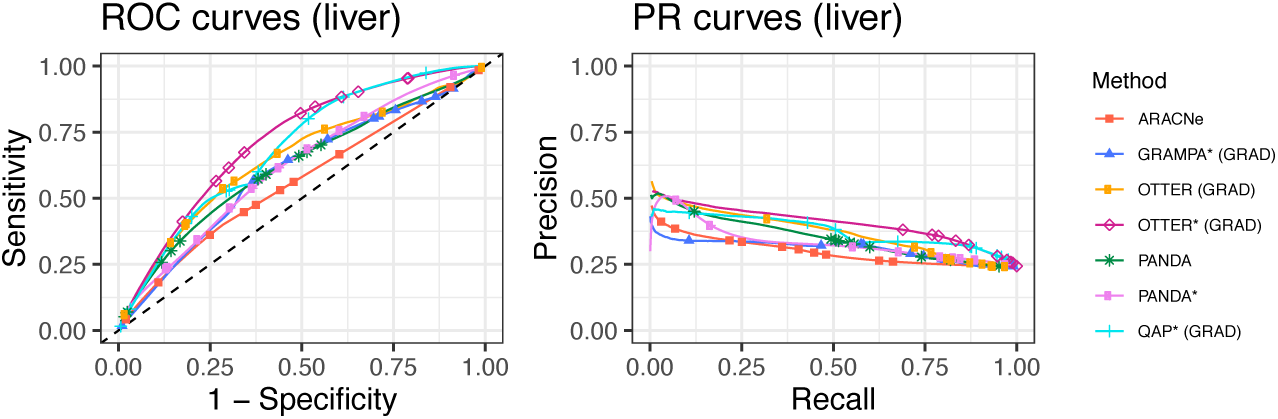
Performance curves for liver tissue.

In general, we observe better performance for the methods that incorporate additional biological evidence such as (transformed) protein-protein interactions and binding motifs, even though these are not tissue specific. A reason for this is that correlations in gene expression can be caused by many factors and that many TFs are expressed at very low levels but strongly activate their target genes, obscuring correlations between TFs and their targets Hence, graph matching approaches are a promising alternative to models that make predictions based on gene expression alone.

## 5 Discussion

We have gained theoretical insights into a state-of-the-art gene regulatory network inference method, PANDA. Our reformulation of the biological intuition behind PANDA as a non-convex optimization problem, OTTER, has multiple global minima, which we have characterized explicitly. Alternative solution approaches can therefore select different minima.

While a gradient descent approach resembles the original PANDA algorithm more closely, the spectral approach selects solutions closest to a biologically reasonable guess based on DNA motif information. The latter has network recovery guarantees in low noise settings and performs well on artificial data, while the former is more robust with respect to noise. The gradient descent outperforms state-of-the art gene regulatory network inference approaches on real world data sets corresponding to three human cancer tissues, which we make publicly available in their processed form for this task.

As we highlight, relaxed graph matching approaches like the relaxed quadratic assignment problem apply to this setting and achieve competitive performance. Hence, we see great potential in transferring alternative relaxed graph matching techniques to gene regulatory network inference in future investigations.

### Data availability

OTTER is available in R, Python, and MATLAB through the netZoo packages: netZooR v0.7 (https://github.com/netZoo/netZooR), netZooPy v0.7 (https://github.com/netZoo/netZooPy), and netZooM v0.5 (https://github.com/netZoo/netZooM).

## Supporting information

proofs and additional information

## Acknowledgements

The results shown here are in part based upon data generated by the TCGA Research Network: https://www.cancer.gov/tcga. DW, MG, CL, JQ, RB were supported by a grant from the US National Cancer Institute (1R35CA220523). JP acknowledges support from the US National Heart, Lung, and Blood Institute (NHLBI): K25HL140186 and KG from the K25 grant: K25HL133599. We thank Alkis Gotovos for helpful feedback on the manuscript.

