## Supplementary material for "Gene Regulatory Network Inference as Relaxed Graph Matching": proofs and additional information

### Supplementary Material: Gene Regulatory Network Inference as Relaxed Graph Matching

**Deborah Weighill**
Department of Biostatistics
Harvard T.H. Chan School of Public Health
Boston, MA 02115


**Marouen Ben Guebila**
Department of Biostatistics
Harvard T.H. Chan School of Public Health
Boston, MA 02115


**Camila Lopes-Ramos**
Department of Biostatistics
Harvard T.H. Chan School of Public Health
Boston, MA 02115


**Kimberly Glass**
Channing Division of Network Medicine,
Brigham and Women's Hospital,
Harvard Medical School,
Department of Biostatistics
Harvard T.H. Chan School of Public Health
Boston, MA 02115


**John Quackenbush**
Department of Biostatistics
Harvard T.H. Chan School of Public Health
Department of Medicine,
Channing Division of Network Medicine,
Brigham and Women's Hospital,
Harvard Medical School,
Boston, MA 02115


**John Platig**
Channing Division of Network Medicine,
Brigham and Women's Hospital,
Harvard Medical School,
Boston, MA 02115


**Rebekka Burkholz**
Department of Biostatistics
Harvard T.H. Chan School of Public Health
Boston, MA 02115


#### 1 Proofs

The central optimization problem to solve is given as

$$\min_{W \in \mathbb{R}^{n_p \times n_p}} f(W) \ \text{ with } \ \mathrm{f(W)} = \frac{(1-\lambda)}{4} \|\mathrm{WW^T - P}\|^2 + \frac{\lambda}{4}\|\mathrm{W^T W - C}\|^2 + \frac{\gamma}{2}\|\mathrm{W}\|^2. \tag{1}$$

##### 1.1 Solutions to the optimization problem

**Theorem 1.** *For given $P \in \mathbb{R}^{n_p \times n_p}$ with $P = P^T$ and $C \in \mathbb{R}^{n_c \times n_c}$ with $C = C^T$, for any spectral decomposition $P = U_p D_p U_p^T$ and $C = V_c D_c V_c^T$, $\lambda \in [0, 1]$, $\Delta := D_c^{[n_p]} - D_p$,*

$$\frac{1}{\lambda^2}\left(\|P\|^2 + \frac{1}{n_p}\operatorname{tr}(P)^2\right) + \frac{1}{n_p}\left(\operatorname{tr}(\Delta)\right)^2 > \|\Delta\|^2, \tag{2}$$

Preprint. Under review.

*the minimization problem (1) has solutions $W^* \in \mathcal{S}$ with singular value decomposition $W^* = U_p D_w V_c^T$, where*

$$d_{w,ii} = \sqrt{\max\left((1-\lambda)d_{p,ii} + \lambda d_{c,ii} - \gamma, 0\right)} \tag{3}$$

*for $i \leq n_p$. For $d_{w,ii} = 0$, the corresponding columns of $U_w$ and $V_w$ are not restricted to eigenvectors of $P$ and $C$. The eigenvalues of $C$ are ordered so that $D_c = D_{c,\pi}$, where the permutation solves the minimization problem*

$$\pi = \mathrm{argmin}_{\pi' \in \mathcal{P}} \frac{\lambda(1-\lambda)}{2} \|\mathrm{D_p} - \mathrm{D}_{\mathrm{c},\pi'}^{[\mathrm{n_p}]}\|^2 - \frac{\lambda}{2}\|\mathrm{D}_{\mathrm{c},\pi'}^{[\mathrm{n_p}]}\|^2 + (1-\lambda)\gamma \,\mathrm{tr}\left(\mathrm{D}_{\mathrm{c},\pi'}^{[\mathrm{n_p}]}\right). \tag{4}$$

***Proof of Theorem 1**.* It is easy to verify that the gradient of the objective

$$f(W) = \frac{(1-\lambda)}{4}\|WW^T - P\|^2 + \frac{\lambda}{4}\|W^TW - C\|^2 + \frac{\gamma}{2}\|W\|^2 \tag{5}$$

is given by

$$\nabla f(W) = WW^TW - (1-\lambda)\frac{1}{2}(P+P^T)W - \frac{1}{2}\lambda W(C+C^T) + \gamma W,$$

which is zero at the extrema of the objective. Note that even for non-symmetric $P$ and $C$, we could solve the problem for the symmetric $(P+P^T)/2$ and $(C+C^T)/2$. Let us rewrite the gradient with respect to a singular value decomposition of $W = U_w D_w V_w^T$

$$0 = \nabla f(W) = U_W\left[D_w D_w^T D_w - (1-\lambda)U_w^T P U_w D_w - \lambda D_w V_w^T C V_w + \gamma D_w\right] V_w^T$$

. Hence, $(1-\lambda)U_w^T(P-\gamma I)U_w D_w + \lambda D_w V_w^T (C - \gamma I)V_w = D$ needs to be a diagonal matrix so that $W$ can be an optimum. Clearly, for $U_w = U_p$ and $V_w = V_c$, we have

$$0 = D_w D_w^T D_w - (1-\lambda)D_p D_w - \lambda D_w D_c + \gamma D_w$$

and solving for $D_w$ leads to $d_{w,ii} = 0$ or $d_{w,ii} = \sqrt{(1-\lambda)d_{p,ii} + \lambda d_{c,ii}}$, where we follow the convention that the singular values of a matrix are positive.

The question remains whether additional zeros of the gradient exist for which $A = (1-\lambda)U_w^T(P - \gamma I)U_w$ and $B = \lambda V_w^T(C - \gamma I)V_w$ are not diagonal matrices but $AD_w + D_w B$ is. As we prove next, this is only possible if $W$ does not have a simple spectrum so that $d_{w,ii} = d_{w,jj}$ for some $j \neq i$.

$W$ is a local minimum iff $AD_w + D_w B = D_w D_w^T D_w$. Hence, $d_{w,ii} = 0$ or $d_{w,ii} = \sqrt{a_{ii} + b_{ii}}$ following the convention that singular values are positive. For off-diagonal elements we have

$$0 = a_{ij} d_{w,ii} + b_{ij} d_{w,jj} \tag{6}$$

Adding the elements corresponding to $ij$ and $ji$ and using the symmetry of $A$ and $B$ we receive $0 = a_{ij}(d_{w,ii} + d_{w,jj}) + b_{ij}(d_{w,ii} + d_{w,jj})$ and thus $a_{ij} = -b_{ij}$ if at least one $d_{w,ii} \neq 0$ or $d_{w,jj} \neq 0$. Plugging this relation back into Eq. (6) leads to $0 = a_{ij}(d_{w,ii} - d_{w,jj})$. For $d_{w,ii} \neq d_{w,jj}$, it follows that $a_{ij} = a_{ji} = 0$ and likewise $b_{ij} = b_{ji} = 0$. Therefore, if $W$ has a simple spectrum with non-zero eigenvalues, $U_w^T U_p$ and $V_w^T V_c$ must be permutation matrices and $W$ of the claimed structure.

So far we have determined the extrema of the objective but which are the global minima? To answer this question, we have to evaluate the objective function, which is $f(W) = (1-\lambda)/4\|P - U_w D_w D_w^T U_w\|^2 + \lambda/4\|C - V_w D_w^T D_w V_w^T\|^2 + \gamma/2\|U_W D_w V_w^T\|^2$ for any $W = U_w D_w V_w^T$. With $\nabla f(W) = 0$ so that $d_{w,ii} = \sqrt{a_{ii} + b_{ii}}$ for $i \leq n_p$, this becomes

$$f(W) = \frac{\lambda(1-\lambda)}{4}\|\tilde{A} - \tilde{B}^{[n_p]}\|^2 + \frac{\lambda}{4}\left(\|C\|^2 - \|D_c^{[n_p]}\|^2\right) + \frac{\gamma}{2}\left(\lambda \,\mathrm{tr}(D_p) + (1-\lambda)\,\mathrm{tr}(D_c^{n_p})\right) - \frac{\gamma^2}{4}n_p, \tag{7}$$

where we define $\tilde{A} = A/(1-\lambda) = U_w^T(P - \gamma I)U_w$ and $\tilde{B} = B/\lambda = V_w^T(C - \gamma I)V_w$ and use properties of the $l_2$ norm and the trace, for instance, that both are invariant under multiplication with orthogonal matrices. Furthermore, we make use of the property $\|M\|^2 =< M, M >$ with respect to the usual l2 scalar product $< M, N >= \sum_{i,j} i', j' m_{ij} n_{i'j'}$ for matrices $M$ and $N$ of the same dimensions. The next step is to simplify

$$\|\tilde{A} - \tilde{B}^{[n_p]}\|^2 = \frac{1}{\lambda^2}\left(\|P - \gamma I\|^2 + \|D_w D_w^T\|^2 - 2 < D_{\tilde{A}}, D_w D_w^T >\right), \tag{8}$$

which is the only part of the objective that distinguishes different minima with the same ordering of $C$'s eigenvalues. We are particularly interested in comparing two cases: (i) $W$ refers to a spectral decomposition of $P$ and $C$ so that $U_w = U_p$ and $V_w = V_c$. (ii) $d_{w,ii} = \operatorname{tr}\left(A + B^{[n_p]}\right)/n_p$ for all $i \leq n_p$, which allows for non-zero off-diagonal elements of $A$ and minimizes $\|D_w D_w^T\|^2$. Note that the following derivations and arguments would also apply to a combination of $(i)$ and $(ii)$, where $A + B^{[n_p]}$ is comprised of different blocks that are of form $(i)$ or $(ii)$ for a subset of the eigenvalues.

In case (i), we have

$$\|\tilde{A}-\tilde{B}^{[n_p]}\|^2 = \|U_w^T U_p(D_p-\gamma I)U_p^T U_w - \left(V_w^T V_c(D_c-\gamma I)V_c^T V_w\right)^{[n_p]}\|^2 = \|D_p - D_c^{[n_p]}\|^2 = \|\Delta\|^2.$$

For case (ii), we utilize representation (8). With $\|D_w D_w^T\|^2 = n_p d_{w,11}^2 = \operatorname{tr}\left(A + B^{[n_p]}\right)^2/n_p = \left(\operatorname{tr}(P)(1-\lambda) + \lambda \operatorname{tr}(D_c^{[n_p]}) - \gamma n_p\right)^2/n_p$ and $< D_{\tilde{A}}, D_w D_w^T >= d_{w,11}^2 < D_{\tilde{A}}, I >= d_{w,11}^2 \operatorname{tr}\left(\tilde{A}\right) = d_{w,11}^2\left(\operatorname{tr}(P) - \gamma n_p\right)$, we obtain

$$\|\tilde{A} - \tilde{B}^{[n_p]}\|^2 = \frac{1}{\lambda^2}\|P\|^2 + \frac{1}{\lambda^2 n_p}\operatorname{tr}(P) + \frac{1}{n_p}\operatorname{tr}(\Delta)^2.$$

In consequence, a global minimum is achieved by (i) in case that $\|\Delta\|^2 < \|\tilde{A} - \tilde{B}^{[n_p]}\|^2 = \frac{1}{\lambda^2}\|P\|^2 + \frac{1}{\lambda^2 n_p}\operatorname{tr}(P) + \frac{1}{n_p}\operatorname{tr}(\Delta)^2$, which proves Condition (2). The optimal matching of eigenvalues of $P$ and $C$ is then given by minimizing $f(W)$, which is for (i):

$$\begin{aligned}
&\min_{\pi} f(W)\\
&= \min_{\pi} \frac{\lambda(1-\lambda)}{4}\|D_p - D_{c,\pi}^{[n_p]}\|^2 + \frac{\lambda}{4}\left(\|C\|^2 - \|D_c^{[n_p]}\|^2\right) + \frac{\gamma}{2}\left(\lambda \operatorname{tr}(D_p) + (1-\lambda)\operatorname{tr}(D_c^{n_p})\right) - \frac{\gamma^2}{4}n_p\\
&= \min_{\pi} \frac{\lambda(1-\lambda)}{2}\|D_p - D_{c,\pi}^{[n_p]}\|^2 - \frac{\lambda}{2}\|D_c^{[n_p]}\|^2 + (1-\lambda)\gamma \operatorname{tr}(D_c^{n_p}),
\end{aligned}$$

where we keep only the terms that are affected by a permutation of the eigenvalues $\pi$. This derives Eq. (4) and concludes the proof. □

#### 1.2 Network recovery

Let $\Phi$ denote the cumulative distribution function (cdf) of a standard normal and $X \sim \mathrm{Ber(p)}$ a Bernoulli random variable with success probability $p$.

**Proposition 2.** *Assume that we observe $P = W^{*T}W^*$, $C = W^* W^{*T}$, and $W_0 = W^* + E$ for a true underlying $W^* \in \mathbb{R}^{n_p \times n_p}$ and noise $E \in \mathbb{R}^{n_p \times n_p}$ with independent identically normally distributed components $e_{ij} \sim \mathcal{N}\left(0, \sigma^2\right)$. Further assume that $P$ and $C$ have a simple spectrum $\{d_1, \ldots, d_{n_p}\}$. Then, for the spectral approach $\widehat{W} = \operatorname{argmin}_{\mathrm{W}\in\mathcal{S}}\|\mathrm{W} - \mathrm{W}_0\|^2$ with $\gamma = 0$, the recovery loss is distributed as $\|\widehat{W} - W^*\|^2 = 4\sum_{i=1}^{n_p} d_i^2 R_i$, where $R_i \sim \mathrm{Ber}\left(\Phi\left(-\mathrm{d_i}/\sigma\right)\right)$ for $d_i > 0$ and $R_i = 0$ for $d_i = 0$ are independent. For any $\epsilon > 0$, it thus holds with the usual Chernoff bound:*

$$\mathbb{P}\left(\|\widehat{W} - W^*\|^2 \leq \epsilon\right) \geq 1 - \exp\left(\epsilon - \mu - \frac{\epsilon}{4}\delta \log\left(\frac{\epsilon}{\mu}\right)\right),$$

*where $\mu = \sum_i p_i$ and $\delta = \frac{1}{\max_i\left(d_{w,ii}^2\right)}$ for $\epsilon \leq \mu$ and $\delta = \frac{1}{\min_i\left(d_{w,ii}^2\right)}$ otherwise.*

***Proof of Proposition 2.*** We assume that the root $W$ is of the form $UD_wD_sV^T$, where we know $U = U_p$, $V = V_c$, and $D_w$ and want to infer the signs $D_s$ with $d_{s,ii} \in \{-1, 1\}$ for $i \leq n_p$. With the input $W_0 = W^* + E$, where the noise $E \in \mathbb{R}^{n_p \times n_p}$ has independent identically normally distributed components $e_{ij} \sim \mathcal{N}\left(0, \sigma^2\right)$, the spectral approach leads to the estimate

$$\widehat{d}_{s,ii} = \operatorname{sign}\left(\sum_{\mathrm{k,l}} \mathrm{u_{ki} w_{0,kl} vli}\right) = \operatorname{sign}\left(\sum_{\mathrm{k,l}} \mathrm{u_{ki}(w *_{kl} + e_{kl}) vli}\right) = \operatorname{sign}\left(\mathrm{d_{w,ii} d^*_{s,ii}} + \sum_{\mathrm{k,l}} \mathrm{u_{ki} e_{kl} vli}\right).$$

To estimate the risk of an error, we therefore need to derive the joint distribution of the random deviations $x_i = \sum_{k,l} u_{ki} e_{kl} vli$. As the $e_{kl} \sim \mathcal{N}\left(0, \sigma^2\right)$ are iid, also their linear combinations $x_i$ are jointly normally distributed with $= 0$ and covariance

$$\mathbb{E}(x_i x_j) = \mathbb{E}\left(\sum_{k,l,k',l'} u_{ki} u_{k'j} vlivl'je_{kl} e_{k'l'}\right) = \sum_{k,k'} u_{ki} u_{k'j} \sum_{l,l'} v_{li} v_{l'j} \mathbb{E}\left(e_{kl} e_{k'l'}\right)$$
$$= \sum_{k,k'} u_{ki} u_{k'j} \sum_{l,l'} v_{li} v_{l'j} \sigma^2 \delta_{k,k'} \delta_{l,l'} = \sigma^2 \sum_{k} u_{ki} u_{kj} \sum_{l} v_{li} v_{lj} = \sigma^2 \delta ij,$$

since $U$ and $V$ are orthogonal and thus also their columns. Thus, $X = (x_1, \ldots, x_{n_p}) \sim \mathcal{N}\left(0, \sigma^2 I\right)$. This allows us to derive the distribution of the sign errors $R_i$, which are defined as $R_i = 1$ if $\widehat{d}_{s,ii} \neq d^*_{s,ii}$ and $R_i = 0$ otherwise. Thus, $R_i = \left(1 - \mathrm{sign}\left(\mathrm{d_{w,ii}} \mathrm{d^*_{s,ii}} + \mathrm{x_i}\right) \mathrm{d^*_{s,ii}}\right)/2$. It follows that these are independent Bernoulli random variables $R_i \sim \mathrm{Ber(p_i)}$ with probability $p_i = \mathbb{P}\left(x_i \leq -d_{w,ii}\right) = \Phi\left(-d_{w,ii}/\sigma\right)$. Consequently, we have a higher error probability for small singular values $d_{w,ii}$.

It is left to show how these sign errors affect the network recovery loss

$$\|\widehat{W} - W^*\|^2 = \sum_i d^2_{w,ii} \left(\widehat{d}_{s,ii} - d^*_{s,ii}\right)^2 = 4 \sum_i d^2_{w,ii} R_i \leq 4 d^2_{w,11} \sum_i R_i,$$

where we assume that the singular values are ordered so that $d^2_{w,ii} \geq d^2_{w,jj}$ for $i < j$. Hence, for all $\epsilon > 0$ we get

$$\mathbb{P}\left(\|\widehat{W} - W^*\|^2 \leq \epsilon\right) = \mathbb{P}\left(\sum_i R_i d^2_{w,ii} \leq \frac{\epsilon}{4}\right) \geq 1 - \min_{t>0} \exp\left(-\epsilon/4t + \mathbb{E}\left(t \sum_i R_i d^2_{w,ii}\right)\right)$$
$$= 1 - \min_{t>0} \exp\left(-\frac{\epsilon}{4} t + \prod_i \left(1 - p_i + p_i e^{d^2_{w,ii} t}\right)\right) \geq 1 - \min_{t>0} \exp\left(-\frac{\epsilon}{4} t + \sum_i p_i (e^{t d^2_{w,ii}} - 1)\right)$$
$$\geq 1 - \exp\left(\epsilon - \mu - \frac{\epsilon}{4} \delta \log\left(\frac{\epsilon}{\mu}\right)\right)$$

for $t = \delta \log\left(\epsilon/\mu\right)$, where $\mu = \sum_i p_i$ and $\delta = \frac{1}{\max_i\left(d^2_{w,ii}\right)}$ for $\epsilon \leq \mu$ and $\delta = \frac{1}{\min_i\left(d^2_{w,ii}\right)}$ otherwise. □

##### 1.3 Gradient dynamics

From a theoretical perspective, we can understand the gradient dynamics for specific choices $W_0$ as initialization. For this purpose, we take the continuous time (for infinitesimally small step size) approximation and study the corresponding gradient flow:

$$\tau \frac{dW}{dt} = -\nabla f(W) = -WW^T W + (1 - \lambda) PW + \lambda WC - \gamma W, \tag{9}$$

where we set the time unit $\tau = 1$ in the following for simplicity. If the initial $W_0$ has a similar singular value decomposition as a solution, the differential equation decouples and we can solve the resulting one-dimensional ordinary differential equations for the diagonal elements explicitly.

**Proposition 3.** *For initial $W_0 = U_p D_0 V_c^T$ with $U_p D_p U_p^T$ and $V_c D_c V_c^T$, the solution of the gradient flow (9) is given by $W(t) = U_p D_t V_c^T$ with*

$$d_{t,ii} = \mathrm{sign(d_{0,ii})} \mathrm{d_{w,ii}} \sqrt{\frac{1}{2} \mathrm{h}\left(\mathrm{d^2_{w,ii}} \mathrm{t} + \mathrm{h}^{-1}\left(2 \frac{\mathrm{d^2_{0,ii}}}{\mathrm{d^2_{w,ii}}} - 1\right)\right) + 1},$$

*where $h(x) = \mathrm{tanh(x)}$ if $d^2_{0,ii} < d^2_{w,ii}$ and $h(x) = \mathrm{coth(x)}$ otherwise.*

***Proof of Proposition 3***. We start from the actual gradient descent, whose updates are discrete in time and given by

$$W(t+1) = W(t) - \eta \nabla f(W)$$

with $W(0) = U_p D_0 V_c^T$ and learning rate $\eta > 0$. We prove inductively that $W(t)$ has singular value decomposition $W(t) = U_p D(t) V_c^T$ (where we also allow for negative singular values). Hence, only the singular values change over time while $U_w = U_p$ and $V_w = V_c$ stay constant. Initially, the induction hypothesis is fulfilled according to our assumption, as $W(0) = U_p D_0 V_c^T$. The induction step assumes that $W(t) = U_p D(t) V_c^T$. Then, $W(t+1) = W(t) - \eta \nabla f(W) = U_p D(t) V_c^T - \eta U_p \left[D(t)D(t)^T D(t) - (1-\lambda) D_p D(t) - \lambda D(t) D_c + \gamma D(t)\right] V_c^T$. Thus, $W(t+1) = U_p D(t+1) V_c^T$ with diagonal $D(t+1) = D(t) - \eta \left[D(t)D(t)^T D(t) - (1-\lambda) D_p D(t) - \lambda D(t) D_c + \gamma D(t)\right]$.

The structure of the differential equation is preserved in the limit of infinitesimal stepsize $\eta$ to obtain the gradient flow:

$$\frac{dD}{dt} = -DD^T D + (1-\lambda) D_p D(t) + \lambda D(t) D_c - \gamma D(t)$$

so that $W(t) = U_p D(t) V_c^T$. Thus, the differential equations are decoupled and the problem is reduced to solving $n_p$ 1-dimensional differential equations of the form $\frac{dx}{dt} = -x^3 + \mu x$, where $\mu$ depends on the respective equation as $\mu_i = (1-\lambda) d_{p,ii} + \lambda d_{c,ii} - \gamma = d_{w,ii}^2$. We can solve this type of equation by rewriting it as

$$x \frac{dx}{dt} = \frac{1}{2} \frac{dx^2}{dt} = -x^4 + \mu x^2.$$

With a change of variable $s = x^2$, this becomes $\frac{ds}{dt} = 2s\left(\mu - s\right)$ so that we can simply integrate

$$\int_{x^2(0)}^{x^2(t)} \frac{1}{\frac{\mu^2}{4} - \left((s - \frac{\mu}{2}\right)^2} ds = 2 \int_0^t dt'.$$

This results in

$$x^2(t) = \frac{\mu}{2} \left[1 + h\left(\mu t + h^{-1}\left(2\frac{x^2(0)}{\mu} - 1\right)\right)\right],$$

where $h = \tanh$ if $x_0^2 < \mu$ and $h = \coth$ otherwise. In both cases, $x^2(t)$ does not pass 0 during its evolution. This is relevant, since we have an ambiguity in the sign of $x(t)$ when we know only $x^2(y)$. Passing through 0 would have been the only option for $x(t)$ to switch signs. Therefore, $x(t) = \mathrm{sign(x(0))}\sqrt{(\mathrm{x^2(t)})}$ inherits the sign of the initial value. Identifying $x(t)$ with $d(t)_i$ and $\mu = d_{w,ii}^2$ concludes the proof. □

#### 2 Correspondence of OTTER to PANDA

PANDA (Passing Attributes between Networks for Data Assimilation) [4] is based on the intuition that a gene regulatory matrix $W$ should be the joint root of the gene-gene interaction matrix $C$ and the protein-protein interaction matrix $P$, i.e. $W^T W \approx P$ and $WW^T \approx P$. This is realized within a message passing framework that iteratively modifies the matrices $C$, $P$, and $W_0$ in discrete time steps $t$ as:

$$P(t+1) = (1-\alpha)P(t) + \alpha \frac{W(t)W^T(t)}{r(W(t), W^T(t))}, \tag{10}$$

$$C(t+1) = (1-\alpha)C(t) + \alpha \frac{W^T(t)W(t)}{r(W^T(t), W(t))}, \tag{11}$$

$$W(t+1) = (1-\alpha)W(t) + \alpha \frac{1}{2}\left(\frac{P(t)W(t)}{r(P(t), W(t))} + \frac{W(t)C(t)}{r(W(t), C(t))}\right), \tag{12}$$

where $r$ denotes a centralization factor $r(M, N) := \sqrt{\|M\|^2 + \|N\|^2 - | < M, N > |}$ that prevents exploding matrix entries and $\alpha \in [0, 1]$ is a tuning parameter that is set to $\alpha = 0.05$ as a default.

In the following, we will discuss how OTTER relates to the main idea of PANDA and ignore the factor $r(\cdot, \cdot)$, as such a scaling is handled differently by the ADAM gradient descent algorithm. As a reminder, the OTTER gradient descent updates are given by

$$W(t+1) = W(t) - \eta\gamma W(t) - \eta W(t) W^T(t) W(t) + \eta(1-\lambda)P(0)W(t) + \eta\lambda W(t)C(0). \tag{13}$$

We claim that these are similar to the PANDA update

$$W(t+1) = (1-\alpha)W(t) + \alpha\frac{1}{2}\left(P(t)W(t) + W(t)C(t)\right). \tag{14}$$

At first glance, we can identify already the first two terms $W(t) - \eta\gamma W(t)$ and $(1-\alpha)W(t)$ for $\alpha = \eta\gamma$. It would be tempting to match also the last two $P(t)W(t) + W(t)C(t)$. Yet, a noticeable difference is that PANDA updates $P(t)$ and $C(t)$ while OTTER keeps them fixed to the input. While we cannot resolve this difference completely, we can capture the dependence of $P(t)$ and $C(t)$ on $W(t)$ more adequately. From Eq. (10) we can deduce $P(t) = \frac{1}{1-\alpha}\left(P(t+1) - \alpha W(t)W^T(t)\right)$ and $C(t) = \frac{1}{1-\alpha}\left(C(t+1) - \alpha W^T(t)W^(t)\right)$ and plug these into Eq. (14):

$$W(t+1) = (1-\alpha)W(t) + \alpha\frac{1}{2}\frac{1}{1-\alpha}\left(P(t+1)W(t) + W(t)C(t+1) - 2\alpha W(t)W^T(t)W(t)\right),$$

which looks almost like our OTTER update (13). Only the time dependence of $P(t+1)$ and $C(t+1)$ cannot be resolved and remains a difference between the two approaches. Yet, considering that $\alpha$ is usually quite small $\alpha \approx 0.05 - 0.1$ and PANDA takes often only $40$ time steps in total, the difference between both approaches is small enough that theoretical insights concerning OTTER should also reflect on PANDA.

### 3 Gene regulation

The central dogma of molecular biology describes the flow of information in a cell from DNA to RNA and finally to proteins. DNA is the blueprint of the functional capacity of a cell, and has certain regions - genes - that can be considered functional units. Each gene codes for a specific protein, which, when produced, performs a specific function in the cell. Gene regions in double-stranded DNA are *transcribed* to a single-stranded mRNA transcript molecule that serves as a template for the protein construction. The mRNA transcript is then *translated* into the corresponding protein. The extent to which a protein is expressed, as measured by protein abundance, thus depends (in part) on the extent to which the gene is expressed (transcribed) and the abundance of the corresponding mRNA. The regulation of genes, including under which conditions and to what degree they are expressed defines a cell's to respond to environmental stimulus, helps define and distinguish individual tissues, allows for developmental processes to occur, and mediates the development and progression of diseases, including their response to theapies. Transcription factors are proteins within the cell that bind to the DNA in "promoter regions" of individual genes and regulate the expression of that gene by recruiting the "transcriptional machinery" to allow mRNA to be synthesized (for a useful review on gene regulation, see [11]). Different transcription factors (TFs) regulate different genes in a many-to-many relationship, and sometimes require co-operativity with each other TFs (Figure S1 A). We can represent the regulatory relationships between TFs and genes as a bipartite network (Figure S1 B). Because the TFs are themselves encoded by genes in the genome, the entire regulatory network represents a complex, adaptive system. Differences between biological states, such as between health and disease, are determined by activation or repression of individual regulatory edges between TFs and genes. Because every cell must carry out some basic functions, including respiration and metabolic processes, much of the network active in any two cells will be identical. It is often the small differences in GRN structure between states that define those states.

### 4 Datasets for gene regulatory network inference

We demonstrate the functionality of OTTER on three cancer datasets from The Cancer Genome Atlas (https://www.cancer.gov/about-nci/organization/ccg/research/structural-genomics/tcga) representing tumors of the liver, cervix, and breast.

#### 4.1 Cancer gene expression data

Gene expression data in The Cancer Genome Atlas (TCGA) [12] was downloaded from recount2 [2] at `https://jhubiostatistics.shinyapps.io/recount/` on 01/10/2020.

##### 4.2 Defining the dimensions of $C$ and $P$

For each tissue, we need to define the dimensions of $C$ and $P$, namely the set of TFs $[n_p] = \{1, \ldots, n_p\}$ and the set of genes $[n_c] = \{1, \ldots, n_c\}$, respectively. A list of all known TF gene names and ENSEMBL ids downloaded `http://humantfs.ccbr.utoronto.ca/download/v_1.01/TFs_Ensembl_v_1.01.txt` and `http://humantfs.ccbr.utoronto.ca/download/v_1.01/TF_names_v_1.01.txt` on 03/09/2020. The reshape R package [2] was used to normalize gene expression measurements as Transcripts Per Million (TPM), which accounts for biases introduced by sample read depth and gene length. For each tissue, we removed genes with consistently low expression, having a TPM $\leq 0.25$ across at least 80% of the samples. This resulted in gene sets of size $n_c = 31,247$ for breast tissue, $n_c = 30,181$ for cervix tissue and $27,081$ for liver tissue.

Transcription factors were found to have, on average, lower expression than other genes (Figures S1-S3), thus, to include a transcription factor in our analysis, we only required that the transcription factor was expressed with a standard deviation $> 0$. All 1,637 were expressed in all tissues, making $n_p = 1,637$ for breast, cervix and liver tissues.

##### 4.3 Constructing $C$, $P$ and $W_0$

###### 4.3.1 Gene co-expression matrix, $C$

For each tissue, the gene co-expression matrix, representing genes likely co-regulated, was constructed by calculating the Pearson correlation coefficient between all pairs of genes. This resulted in the square matrix $C$, with dimension $n_c = 31,247$ for breast tissue, $n_c = 30,181$ for cervix tissue and $27,081$ for liver tissue.

###### 4.3.2 Protein-protein interaction matrix, $P$

A list of protein-protein interactions used in [10] was downloaded from `https://sites.google.com/a/channing.harvard.edu/kimberlyglass/tools/gtex-networks` on 09/09/2019. These protein-protein interactions were filtered to those involving transcription factors, and used to populate the $n_p \times n_p$ PPI matrices $P$. Pairs of TFs for which no interaction data was available were set to 0.

###### 4.3.3 Motif-based GRN prior, $W_0$

The initial estimate $W_0$ of the gene regulatory network is constructed based on TF motif information. The FIMO software [5] from MEME suite [1] was used to scan the hg38 human genome assembly for known TF motifs - sequences which are predicted to be bound by specific TFs. Motif matches with a p-value $\leq 10^{-4}$ were considered significant. The positions and IDs of Ensembl genes in the hg38 genome assembly were downloaded from the UCSC Table browser `https://genome.ucsc.edu/cgi-bin/hgTables` on 11/13/2019, and from `https://www.gencodegenes.org/human/` on 03/09/2020. Gene promoter regions were defined as the 1000bp (base pair) region [-750bp, +250bp] around the gene's annotated transcriptional start site, taking into account the strand on which the gene resides. The GenomicRanges R package [7] was used to overlap motif hits with gene promoters, resulting in a set of TF-gene associations, indicating that a motif of a TF was found overlapping with the promoter region of a gene. These associations were then used to populate the $n_p x n_c$ matrix $W_0$ with $W_0[ij] = 1$ if the motif of TF $i$ was found in the promoter of gene $j$ and $W_0[ij] = 0$ otherwise.

##### 4.4 ChIP-seq experimental data for validation

Chromatin immunoprecipitation (ChIP)-seq is a technique that allows for the experimental identification of protein-DNA interactions, and can thus provide a validation data set for TF binding of gene promoters. ChIP-seq experiments are performed individually per TF, and thus, one can only obtain binding site information for TFs that have specifically been studied before with this technique. ChIP-seq data consisting of the genome-wide binding regions of select TFs for the HeLa cervical cancer cell line (48 TFs), HepG2 liver cancer cell line (77 TFs) and MCF7 breast cancer cell line (62 TFs) were downloaded from the ReMap2018 database `http://pedagogix-tagc.univ-mrs.fr/remap/` on 01/15/2020. These cell lines represent the closest tissues to those of the expression data for which

ChIP-seq data was available. We recognize that many cancers have distinct subtypes that often differ substantially from one-another; we ignore those subtypes given the limitations of the available data and recognizing that subtype differences will be smaller than differences between cervix, liver, and breast tumors ChIP-seq-determined binding regions for each TF in each cell line were mapped to the promoter regions of genes in the same manner as described for motif regions. This allowed us to construct a validation regulatory subnetwork, allowing us to validate the our predicted regulatory relationships from OTTER (and other GRN estimation methods) for the portion of TFs for which ChIP-seq data was available. Precision-recall and ROC curves were calculated using the precrec R package [9].

### 5 Algorithms and model parameters

The best performing methods are based on ADAM gradient descent [6] as stated below.

**Algorithm 1: ADAM gradient descent.**

**Inputs:** $W$, $P$, $C$, $\nabla f(\cdot)$
**Parameters:** $\eta$ (learning rate), $I$ (number of iterations)
$\beta_1 = 0.9$; $\beta_2 = 0.999$; $\epsilon = 0.00000001$;
$\beta_{1,t} = \beta_1$; $\beta_{2,t} = \beta_2$;
$m = 0$; $v = 0$;
**for** $i = 1, \ldots, I$ **do**
  $m = \beta_1 m + (1 - \beta_1)\nabla f(W, P, C)$;
  $v = \beta_2 v + (1 - \beta_2)(\nabla f(W, P, C))^2$;
  $\beta_{1,t} = \beta_{1,t}\beta_1$; $\beta_{2,t} = \beta_{2,t}\beta_1$;
  $\alpha = \eta \frac{\sqrt{1-\beta_{2,t}}}{1-\beta_{1,t}}$;
  $\epsilon_t = \epsilon\sqrt{1 - \beta_{2,t}}$;
  $W = W - \alpha \frac{m}{\epsilon_t + \sqrt{v}}$;
**end for**

We adapt the learning rate $\eta$, exponent $b$, the number of iterations $I$ and the parameters of the gradient for each method that can also take different transformations of the original matrices $P_0$, $C_0$, $W_0$ into account. Our specific choices are listed in Table S1. The respective gradients are defined as $\nabla f = 4WW^TW - 4(1-\lambda)PW - 4\lambda WC + 2\gamma W$ for OTTER, $\nabla f = P^2W + WC^2 - 2PWC + \gamma W$ for QAP, and $\nabla f = P^2W + WC^2 - 2PWC + \gamma W - \delta J$ for GRAMPA. We added another version of OTTER gradient descent (OTTER$^*$ grad 2) to the table with a different scaling of the protein-protein interaction matrix $P$. This version performs slightly worse than the reported alternative OTTER$^*$ grad, but has the advantage that the gene regulatory network inference improves constantly with every gradient step. All other approaches, i.e. OTTER$^*$ grad, GRAMPA$^*$ grad, QAP$^*$ grad, and the non-transformed counterparts, cycle through phases of excellent performance and very reduced performance making them less robust to the wrong choice of $I$. The remaining methods are parameterized as follows. OTTER (spectral) uses the parameters $\mu = 0.0043208$ and $\lambda = 0.99498$, while the one based on transformed inputs uses $\mu = 0.335$ and $\lambda = 0.0035$. GENIE3 and TIGRESS have been computed only with respect to transcription factors that meet the gene expression filter criterion. Accordingly, we have also restricted the validation to those transcription factors. This was meant to make the task easier but the algorithms still showed inferior performance with respect to binding prediction. We used the default parameters, i.e. random forests consisting of 1000 trees and $K = \sqrt{p}$ for GENIE3 and $nstepsLARS = 5$, $\alpha = 0.2$, and $nsplit = 100$ for TIGRESS. Furthermore, we had to restrict TIGRESS and PAR COR to the much smaller number of transcription factors in our validation set to reduce the computational load.

### 6 Biological validation

In order to determine whether OTTER networks capture expected biological functional information, we constructed an OTTER network representing healthy liver and cancerous liver, and investigated the biological functions enriched in areas of the networks which differ between cancer vs. healthy expression data.

A similar process as described above was used to construct process gene expression data and construct OTTER networks. The healthy OTTER network was constructed making use of liver gene expression data from The Genotype Tissue Expression Project (GTEx) [8], whereas the cancer network was constructed using expression data from TCGA [12]. Both gene expression datasets were downloaded from recount2 [12].

From the resulting OTTER networks, a difference network was constructed by taking the absolute value of the difference between corresponding edge weights of the two networks. Lastly, we calculated TF degrees and gene degrees in the resulting difference network, allowing genes and TFs to be ranked by these degrees. Genes/TFs with high "difference degree" are thus those whose neighboring edges have large differences between healthy and cancer networks.

GOrilla [3] was then used to determine which Gene Ontology (GO) biological process terms were enriched in the top of the ranked lists vs the bottom, thus determining which biological functions are enriched in those genes/TFs. Enrichment results (Figures S7 and S8, Tables S2, S3) highlighted several expected cancer-related functions and pathways, including cell proliferation, cell differentiation, cell death, cell-cell adhesion, cell motility, immune-related processes, cellular response to growth factors. OTTER networks thus capture expected biological signal related to the context of the gene expression data.

#### 7 Supplementary Figures and Tables

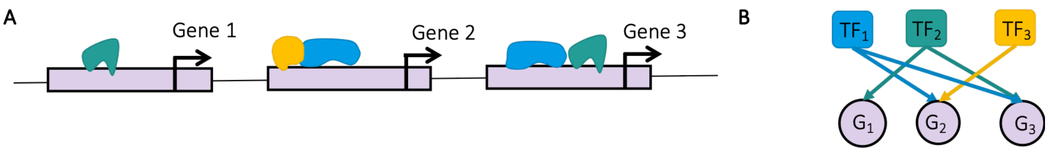


Figure S1: Gene regulation. A. Transcription factors (TFs) are represented by green, blue, and yellow objects that bind to the genome (gray band) in vicinity of the start site of a gene (black error) to regulate its expression. B. Representation of A as bipartite gene regulatory network.

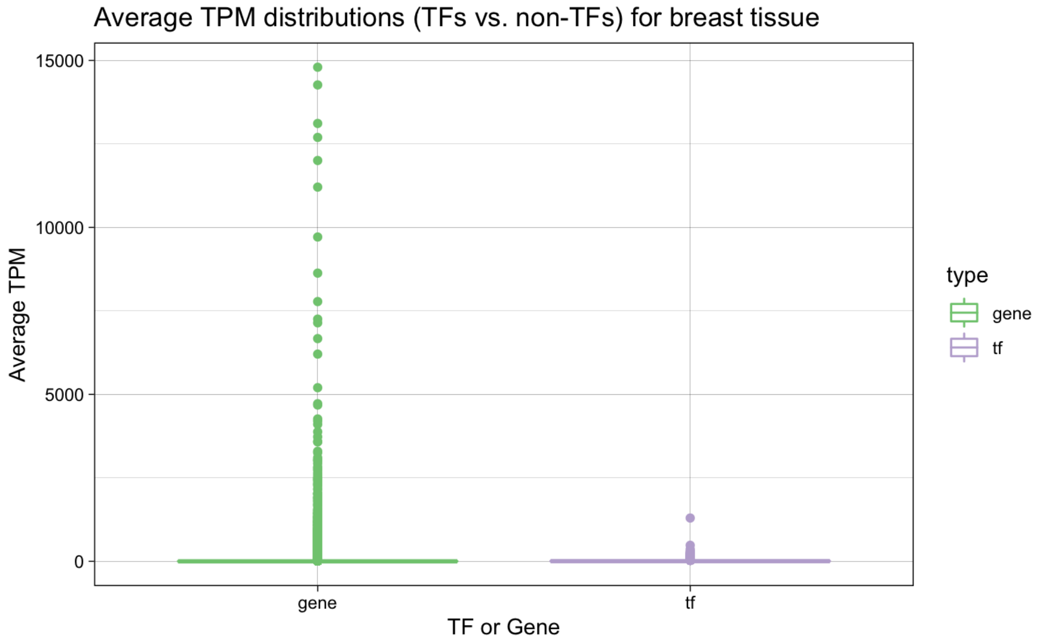


Figure S2: TPM distributions for breast tissue.

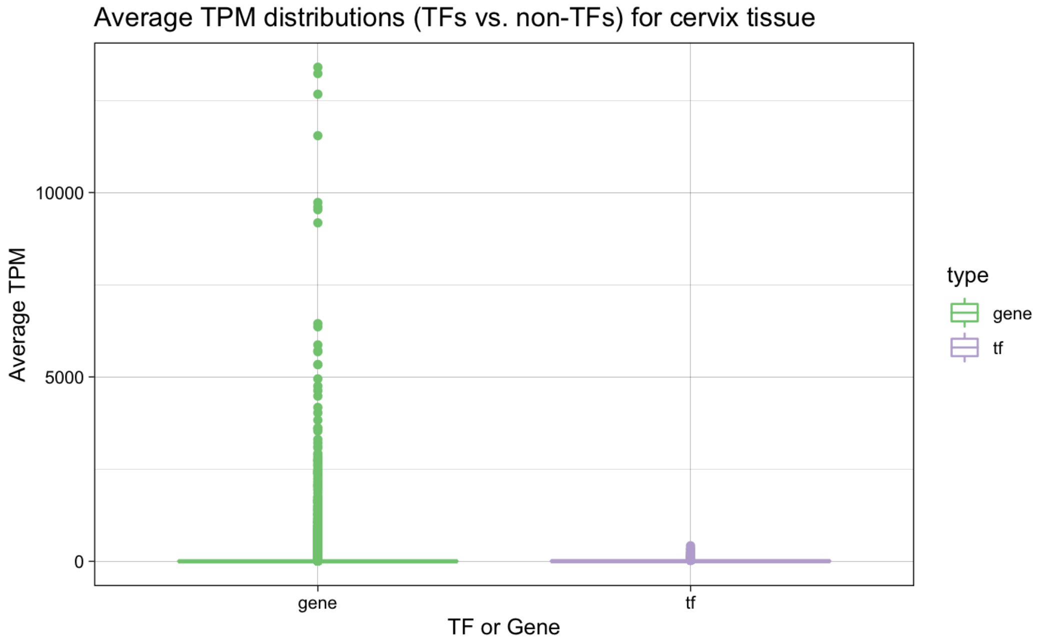


Figure S3: TPM distributions for cervix tissue.

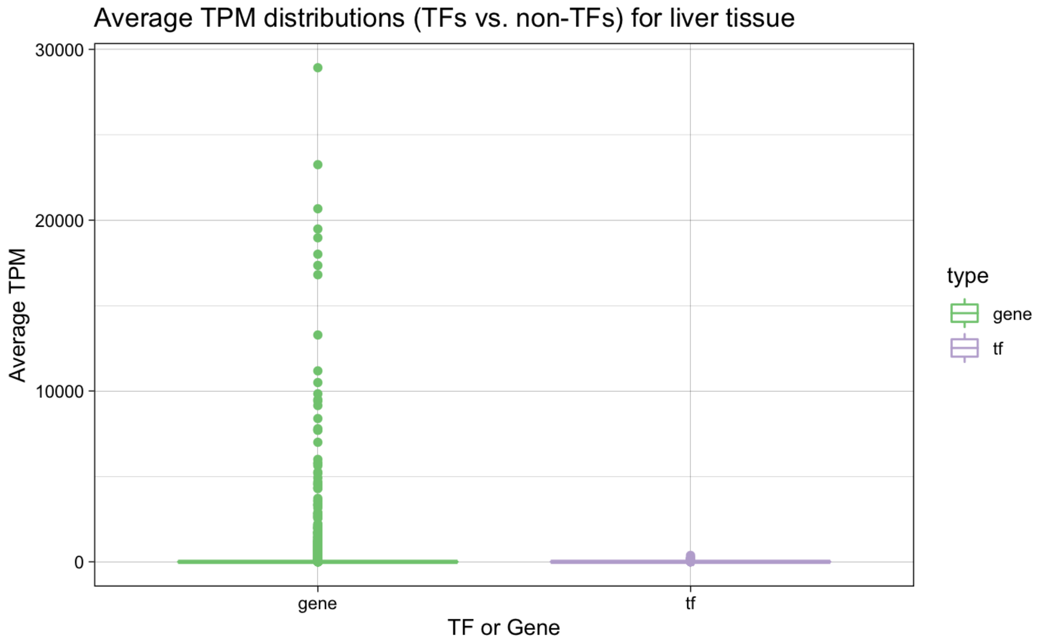


Figure S4: TPM distributions for liver tissue.

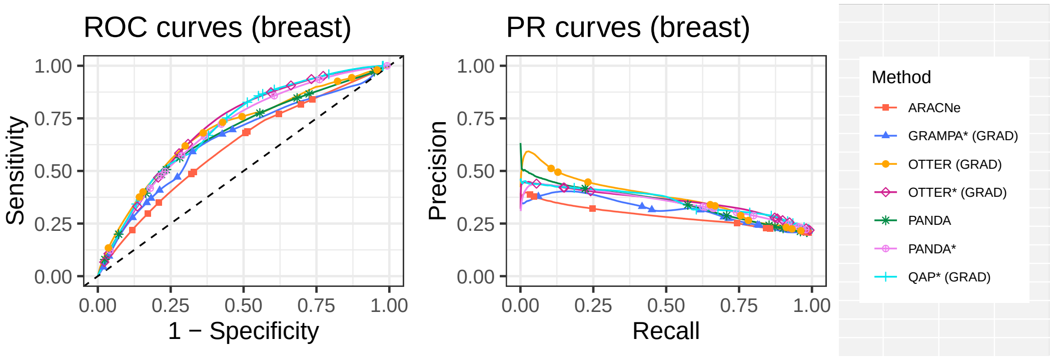


Figure S5: Performance curves for breast.

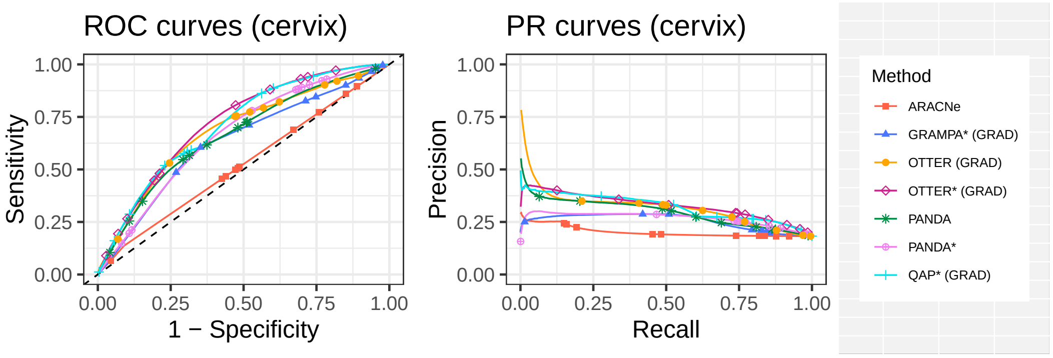


Figure S6: Performance curves for cervix.

Figure S7: Gene Ontology (GO) term enrichment visualization for healthy (GTEx) vs cancerous (TCGA) liver tissue gene differential degree. Enrichment performed and visualized using GOrilla [3]. *See attached file* `FigureS7.png`

Figure S8: Gene Ontology (GO) term enrichment results for healthy (GTEx) vs cancerous (TCGA) liver tissue TF differential degree. Enrichment performed using GOrilla [3]. *See attached file* `FigureS7.png`

Table S1: Parameters for models taking $P$ and $W_0$ into account.

| | Inputs | | | Parameters | | | |
|---|---|---|---|---|---|---|---|
| | $W$ | $P$ | $C$ | $\eta$ | $I$ | $\gamma$ | misc. |
| GRAMPA grad | $W_0/\operatorname{tr}(W_0W_0^T)$ | $P_0/\operatorname{tr}(P_0)$ | $C_0/\operatorname{tr}(C_0)$ | 0.01 | 6 | 0.05 | $\delta = 1$ |
| QAP grad | $W_0/\operatorname{tr}(W_0W_0^T)$ | $P_0/\operatorname{tr}(P_0)$ | $C_0/\operatorname{tr}(C_0)$ | 0.01 | 12 | 0.5 | - |
| OTTER grad | $W_0/\sqrt{\operatorname{tr}(W_0W_0^T)}$ | $P_0/\operatorname{tr}(P_0)$ | $C_0/\operatorname{tr}(C_0)$ | 0.00001 | 60 | 0.335 | $\lambda = 0.035$ |
| PANDA | $W_0$ | $P_0$ | $C_0$ | - | 40 | - | $\alpha = 0.1$ |
| | Transformed $P$ and $W_0$ | | | | | | |
| GRAMPA* grad | $W_0/\sqrt{\operatorname{tr}(W_0W_0^T)}$ | $(P_0+0.05)/\operatorname{tr}(P_0+0.05)$ | $C_0/\operatorname{tr}(C_0)$ | 0.01 | 13 | 0.05 | $\delta = 1$ |
| QAP* grad | $W_0/\sqrt{\operatorname{tr}(W_0W_0^T)}$ | $(P_0+0.05)/\operatorname{tr}(P_0+0.05)$ | $C_0/\operatorname{tr}(C_0)$ | 0.01 | 22 | 0.05 | - |
| OTTER* grad 2 | $(P_0+2.2)W_0/\sqrt{\operatorname{tr}((P_0+2.2)W_0W_0^T(P_0+2.2)^T)}$ | $(P_0+2.2)/\operatorname{tr}(P_0)$ | $C_0/\operatorname{tr}(C_0)$ | 0.00001 | 300 | 0.335 | $\lambda = 0.0035$ |
| OTTER* grad | $(P_0+2.2)W_0/\operatorname{tr}((P_0+2.2)W_0W_0^T(P_0+2.2)^T)$ | $(P_0+2.2)/\operatorname{tr}(P_0+2.2)$ | $C_0/\operatorname{tr}(C_0)$ | 0.00001 | 32 | 0.335 | $\lambda = 0.0035$ |
| PANDA* | $(P_0+2.2)W_0$ | $P_0+2.2$ | $C_0$ | - | ca. 40 | - | $\alpha = 0.1$ |

Table S2: Gene Ontology (GO) term enrichment results for healthy (GTEx) vs cancerous (TCGA) liver tissue gene differential degree. Enrichment performed using GOrilla [3]. *See attached file* `TableS2.xlsx`

Table S3: Gene Ontology (GO) term enrichment results for healthy (GTEx) vs cancerous (TCGA) liver tissue TF differential degree. Enrichment performed using GOrilla [3]. *See attached file* `TableS3.xlsx`

#### 8 Supplementary Data

**Data S1: Breast** The complete set of matrices used to run and validate regulatory networks in breast tissue, including the PPI prior, motif prior, expression matrix, gene and TF names, as well as true positive ChIP-seq edges for validation.

**Data S2: Cervix** The complete set of matrices used to run and validate regulatory networks in cervix tissue, including the PPI prior, motif prior, expression matrix, gene and TF names, as well as true positive ChIP-seq edges for validation.

**Data S3: Liver** The complete set of matrices used to run and validate regulatory networks in liver tissue, including the PPI prior, motif prior, expression matrix, gene and TF names, as well as true positive ChIP-seq edges for validation.

**Data S4: Regulatory Network Outputs** Regulatory networks constructed using the different methods described in the main manuscript.

**Data S5: Script Set 1** Set of scripts required to create the data matrices, as well as run PANDA and ARACNe. This dataset also contains the inputs and outputs required/produced by these scripts.

**Data S6: Data in Matlab** The Matlab file data.mat contains the training and test data for all three tissues for convenient use in Matlab.

#### 9 Supplementary File Links

**README:**
`https://ottersm.s3.amazonaws.com/README.txt`

**Supplementary Figures:**
`https://ottersm.s3.amazonaws.com/OTTER_Supplementary_Figures.tgz`

**Supplementary Tables:**
`https://ottersm.s3.amazonaws.com/OTTER_Supplementary_Tables.tgz`

**Supplementary Data:**
`https://ottersm.s3.amazonaws.com/OTTER_Supplementary_Data.tgz`

[6] Diederik Kingma and Jimmy Ba. Adam: A method for stochastic optimization. *International Conference on Learning Representations*, 12 2014.

[12] Katarzyna Tomczak, Patrycja Czerwińska, and Maciej Wiznerowicz. The Cancer Genome Atlas (TCGA): an immeasurable source of knowledge. *Contemporary oncology*, 19(1A):A68, 2015.
